# A refined single cell landscape of haematopoiesis in the mouse foetal liver

**DOI:** 10.1101/2023.03.01.530564

**Authors:** Elena Ceccacci, Emanuela Villa, Fabio Santoro, Saverio Minucci, Christiana Ruhrberg, Alessandro Fantin

**Affiliations:** Department of Experimental Oncology, IEO, European Institute of Oncology IRCCS, Milan, Italy; University of Milan, Department of Biosciences, Via G. Celoria 26, 20133, Milan, Italy; University of Milan, Department of Oncology and Hemato-Oncology, Via Santa Sofia 9, 20122, Milan, Italy; UCL Institute of Ophthalmology, University College London, 11-43 Bath Street, London EC1V 9EL, UK

## Abstract

During prenatal life, the foetal liver is colonised by several waves of haematopoietic stem and progenitor cells (HSPCs) to act as the main haematopoietic organ. Single cell (sc) RNA-seq has been used to identify foetal liver cell types via their transcriptomic signature and to compare gene expression pattern as haematopoietic development proceeds. To obtain a refined single cell landscape of haematopoiesis in the foetal liver, we have generated a novel scRNA-seq dataset from whole mouse E12.5 liver that includes a larger number of cells than prior datasets at this stage and was obtained without cell type preselection to include all liver cell populations. We combined mining of this dataset with that of previously published datasets at other developmental stages to follow transcriptional dynamics as well as cell cycle state of developing haematopoietic lineages. Our findings corroborate several prior reports on the timing of liver colonisation by HSPCs and the emergence of differentiated lineages and provide further molecular characterisation of each cell population. Extending these findings, we demonstrate the existence of a foetal intermediate haemoglobin profile in the mouse, similar to that previously identified in humans, and a previously unidentified population of primitive erythroid cells in the foetal liver.

## Introduction

The liver is a metabolic hub that regulates glucose and lipid metabolism as well as protein and bile synthesis, but it is also an essential site for both blood and immune system development in mammals. Thus, the foetal liver provides a suitable microenvironment for the expansion and maturation of several waves of multipotent haematopoietic progenitors (Barone et al., 2022; Canu and Ruhrberg, 2021). At the early Carnegie Stage (CS) 10 in humans, corresponding to embryonic day (E) 8.75 in mouse, the liver rudiment emerges as a diverticulum from the ventral domain of the embryonic foregut. Between E9 and E10 in the mouse, the diverticulum transitions from a monolayer of cuboidal endoderm cells into a pseudostratified multilayer of hepatoblasts to form the liver bud (Gordillo et al., 2015). The hepatoblasts then serve as bi-potent progenitors to produce the two major epithelial cell types of the adult liver, hepatocytes and cholangiocytes (also known as biliary epithelial cells), and form the foetal liver (Gordillo et al., 2015). Hepatoblast specification and expansion requires endothelial cells, which arise adjacent to the mouse liver diverticulum at E9.0 (Gordillo et al., 2015) and specialise to form the lining of the hepatic sinusoids as so-called liver sinusoidal endothelial cells (LSECs). In each hepatic acinus, liver sinusoids are sandwiched between cords of hepatocytes to transport blood from the portal veins and hepatic arteries towards draining central veins, with adult human LSECs showing zone-specific heterogeneity (MacParland et al., 2018; Strauss et al., 2017).

Before the bone marrow is established shortly before birth as the main postnatal haematopoietic organ, the liver bud is colonized by different waves of haematopoietic stem and progenitor cells (HSPCs) (Barone et al., 2022; Canu and Ruhrberg, 2021). These include EMPs, which emerge from the yolk sac endothelium from E8.5 onwards in the mouse, and HSPCs originating from E10.5 onwards in the intra-embryonic aorta-gonad-mesonephros (AGM) region, including the first definitive HSCs (Ginhoux and Guilliams, 2016; Mass et al., 2016; McGrath et al., 2015b). Recent pulse chase lineage tracing studies in the mouse suggested that HSCs do not contribute significantly to the foetal erythroid, megakaryocyte and myeloid lineages until late in gestation (Gomez Perdiguero et al., 2015; Soares-da-Silva et al., 2021; Yokomizo et al., 2022).

In adult mammals, erythrocyte differentiation starts when megakaryocyte-erythroid progenitors (MEPs) arising from definitive HSCs progressively differentiate into lineage committed erythroid progenitors, erythroid burst forming unit (BFU-E), erythroid colony forming unit (CFU-E), nucleated proerythroblast, basophilic, polychromatophilic and orthochromatic erythroblast stages, followed by enucleation and formation of reticulocytes and then mature erythrocytes (Chen et al., 2009; Dzierzak and Philipsen, 2013). As opposed to the formation of enucleated erythrocytes, the first erythroid cells during embryonic development arise from primitive haematopoietic progenitors in the extra-embryonic yolk sac blood islands at around CS 7-8 (16-18.5 dpc) in the human (Ivanovs et al., 2017) and E7.5-8.0 in the mouse (Palis et al., 1999). These primitive erythroid cells retain their nucleus and circulate into embryonic organs, including into the liver, where they interact with macrophages in erythroblastic islands to undergo enucleation between E12.5 and E14.5 (Dzierzak and Philipsen, 2013). After the primitive erythroid wave first arises in the yolk sac, a transient definitive wave of yolk sac-derived EMPs produces CD131 (*Csf2rb*)-positive megakaryocyte-erythroid progenitors (MEPs), which seed the foetal liver to provide the main source of erythrocytes and megakaryocytes in the late gestation embryo (Dzierzak and Philipsen, 2013; Iturri et al., 2021; Soares-da-Silva et al., 2021). Transient definitive erythroblasts also interact with liver macrophages to expel their nucleus before entering the circulation at the reticulocyte stage (Dzierzak and Philipsen, 2013).

Erythrocytes express large amounts of haemoglobin, whose subtypes change during prenatal development. Specifically, oxygen affinity decreases from embryonic to foetal to adult haemoglobin, which is thought to accommodate the changing oxygen tension as the embryo develops to maturity (Dzierzak and Philipsen, 2013; Sankaran and Orkin, 2013). All haemoglobins are tetramers composed of two α-like and two β-like globin chains (Sankaran and Orkin, 2013). This globin nomenclature derives from the two α and β chains that are present in the main adult human form. In adult humans, the most common form is α2β2, in which the two α and β subunits are encoded by the *HBA1* or *HBA2* and *HBB* genes, respectively; the rarer tetramer α2δ2 is instead encoded by *HBA1* or *HBA2* and *HBD*. The human embryonic haemoglobins include the following tetramers: Gower-1 ζ2ε2 (*HBZ, HBE1*), Gower-2 α2ε2 (*HBA1/2, HBE1*), Portland-1 ζ2γ2 (*HBZ*, *HBG1* or *HBG2*) and Portland-2 ζ2β2 (*HBZ, HBB*) (Huret et al., 2013). The human foetal haemoglobins (HbF) are termed α2γ2 (encoded by *HBA1* or *HBA2* and *HBG1* or *HBG2*) (Huret et al., 2013); these haemoglobins are thought to be a specific feature of anthropoid primates (Dzierzak and Philipsen, 2013). In the mouse, embryonic β-like globins (εy and βH1) are thought to be restricted to primitive erythrocytes, whereas adult β globin chains are already present in foetal liver-derived erythrocytes (Trimborn et al., 1999). Whether transient definitive erythrocytes in the mouse foetal liver have unique haemoglobin profiles that differ from those of primitive and definitive erythrocytes is not completely understood.

EMPs and other HSPCs in the foetal liver also contribute to myeloid cell production. In particular, EMPs generate liver monocytes that differentiate into tissue-resident macrophages in many organs (Hoeffel and Ginhoux, 2018). These liver monocyte-derived macrophages gradually replace the initial pool of tissue macrophages that is derived from earlier yolk sac primitive progenitors, except for the brain, where microglia derived from primitive progenitors persist (Ginhoux and Guilliams, 2016; Gomez Perdiguero et al., 2015; Hoeffel et al., 2015; Hoeffel and Ginhoux, 2018; Schulz et al., 2012). Other foetal liver myeloid lineages include granulocytes and mast cells (McGrath et al., 2015a). However, a comprehensive single cell transcriptomic profiling of these early myeloid populations has not been described yet.

Given the major contribution to both erythropoiesis and myelopoiesis, investigating the molecular and cellular landscape of early liver development has the promise to increase our understanding the causes of congenital immunodeficiencies, anaemia and also childhood leukaemia (Cazzola et al., 2020). Towards this aim, several single cell (sc) RNA-seq datasets have been recently generated, which sought to identify foetal liver cell types via their transcriptomic signature and compared gene expression patterns at single cell level across the cell types in the foetal liver (e.g., (Dong et al., 2018; Gao et al., 2022; Su et al., 2017; Wang et al., 2020)). However, the haematopoiesis studies using foetal liver scRNA-seq to date focussed on the analysis of selected cell subsets, isolated by genetic lineage-tracing (Freyer et al., 2020) or surface phenotyping (Gao et al., 2022). Other scRNA-seq datasets were generated without isolation bias, but from only a small number of foetal liver cells (Dong et al., 2018; Su et al., 2017), which is suboptimal for deep phenotyping, or they were comprised of larger cell numbers but analysed mostly hepatocyte development (Wang et al., 2020). Here we describe the generation of a scRNA-seq dataset from E12.5 mouse foetal liver, which was obtained without cell preselection and includes a larger number of cells than prior datasets at this stage. We have combined the analysis of our dataset with data mining of other publicly available datasets to provide new insights into early haematopoietic development in the foetal liver and also included an analysis of foetal liver-constituent cell types.

## Materials and methods

### Library construction with 10x Genomics platform

The E12.5 liver scRNA-seq dataset was generated from C57BL/6J foetal liver. Briefly, a single cell suspension from one E12.5 liver, was mechanically and enzymatically homogenized in RPMI1640 with 2.5% foetal bovine serum (FBS, ThermoFisher), 100 μg/ml collagenase/dispase (Roche), 50 μg/ml DNase (Qiagen) and 100 μg/ml heparin (Sigma), followed by purification from debris, dead cells and doublets through fluorescence activated cell sorting (FACS) (**Fig. 1a**). To maximise the yield in droplet encapsulation of single cells, two separate technical replicates of the sample were independently processed.cDNA Droplet-based digital 3’ end scRNA-seq was performed on a Chromium Single-Cell Controller (10X Genomics, Pleasanton, CA) using the Chromium Single Cell 3’ Reagent Kit v2 according to the manufacturer’s instructions. Cells were divided into 2 samples and independently partitioned in Gel Bead-in-EMulsion (GEMs) droplets and lysed, followed by RNA barcoding, reverse transcription and PCR amplification (12–14 cycles). Sequencing-ready scRNA-seq libraries were prepared according to the manufacturer’s instructions, checked and quantified on TapeStation 2200 (Agilent Genomics, Santa Clara, CA) and Qubit 3.0 (Invitrogen, Carlsbad, CA) instruments. Sequencing was performed on a NovaSeq 6000 machine (Illumina, San Diego, CA) using the NextSeq 500/550 High Output v2 kit (75 cycles), at the genomic facility at the European Institute of Oncology (IEO, Italy).

**Figure 1.**
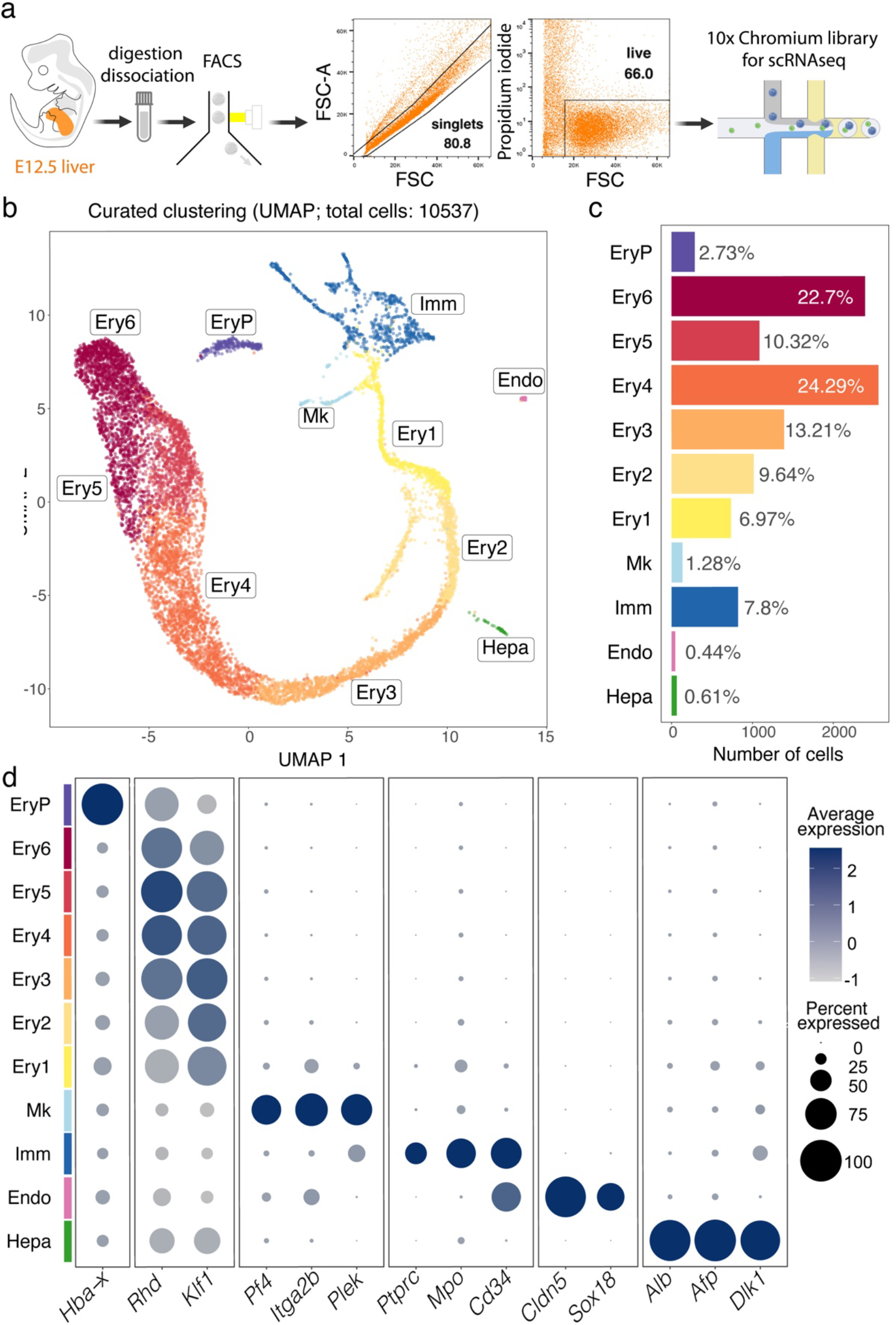
scRNA-seq analysis of the E12.5 mouse liver. (**a**) Gating strategy for E12.5 liver cell isolation prior to single cell library preparation. (**b-d**) A UMAP plot shows distinct cell types (**b**), a histogram shows the percentage and number of cells in each cluster (**c**), and a bubble plot shows expression of selected marker genes in each cluster (**d**). Dot size in (**d**) represents the percentage of cells within a cluster in which a marker was detected, dot colour intensity represents the average expression level of that marker. Abbreviations: Endo, endothelial cells; Ery1-6, erythroid cells; EryP, primitive erythrocytes; Hepa, hepatoblasts; Imm, immune cells; Mk, megakaryoblasts.

### scRNA-seq processing and analysis

Fastq.gz files were generated from raw Illumina BCL files using 10X Genomics Cell Ranger version 2.1.1 with default parameters. The quality of sequencing reads was evaluated using FastQC v0.11.5 and MultiQC v1.5. Cell Ranger v2.1.1 was then used to align the sequencing reads to the mm10 mouse transcriptome and to quantify the expression of transcripts in each cell. Only confidently mapped reads, non-PCR duplicates, with valid barcodes and UMIs were retained to compute a gene expression matrix containing the number of UMI for every cell and gene. All downstream analyses were implemented using R v3.6.1 and the package Seurat v3.1.1 (Butler et al., 2018; Stuart et al., 2019). For each sample, cells expressing less than 300 unique genes and genes expressed in less than 3 cells were discarded. Raw counts were then merged and normalized using the Seurat function SCTransform after removing cells having more than 4% mitochondrial genes. During the normalization, the confounding sources of variation i.e., the mitochondrial mapping percentage, the Sand G2M scores and the difference between them were removed. Cell cycle scores were calculated using CellCycleScoring function. The number of variable features in the SCTransform function after ranking by residual variance was set to 1000. During the normalization, 819 genes were automatically removed because expressed in less than 3 cells independently from the overall counts leading to retain 14,817 total genes. Expression data were than scaled using ScaleData function regressing on S and G2M scores. Graphbased clustering: most variable genes across the merged dataset (i.e., the highly variable genes) were identified based on the highest standardized variance. The procedure is implemented in the FindVariableFeatures function with method = ‘‘vst.” A total number of 1000 genes was selected as top variable features and used to perform the PCA dimensionality reduction. A KNN graph based on the euclidean distance in the 50 PCs space was constructed using the FindNeighbors function.

## Results

### Major cell populations in E12.5 mouse liver in a novel scRNA-seq dataset

To define the cellular landscape of early liver haematopoiesis in the context of other developing liver cell lineages, we generated a scRNA-seq dataset of the mouse E12.5 liver using a droplet-based approach that allowed recovery of a hundred times higher number of cells compared to a previous report, which only sequenced ~100 cells at this stage (Su et al., 2017). Thus, two independent single cell cDNA libraries were prepared from the total single cell suspension of E12.5 foetal liver and sequenced in two batches that were pooled to increase the number of cells sequenced (**Fig. 1a**; **Fig. S1a**). The data from 10,537 cells passed quality control and underwent graph-based unsupervised Louvain clustering and dimensionality reduction by Uniform Manifold Approximation and Projection (UMAP) (**Fig. S1b**). Differentially expressed genes (DEGs, **Fig. S1c**) were used to annotate cell clusters and to pool the most similar clusters into a curated representation of E12.5 liver cell populations (**Fig. 1b-d**). These DEGs are listed in detail in the sections below, where we will discuss specific cell populations. Overall, we identified 11 major cell populations (**Fig. 1b,c**). The identity of each cluster was assigned by matching the cluster expression profile to established cell type-specific genes for erythroid cells (e.g. *Hba-x* for EryP, *Rhd* and *Klf1* for Ery1-6), megakaryocytes (Mk, e.g. *Pf4, Itga2b* and *Plek*), immune cells (Imm, e.g. *Ptprc* and *Mpo*), endothelial cells (Endo, e.g. *Cldn5* and *Sox18*) and hepatocytes (Hepa, e.g. *Alb, Alf* and *Dlk1*) (**Fig. 1d**). As expected, given the red colour appearance of the foetal liver, ~90% of all E12.5 liver cells were erythroid cells (**Fig. 1c**). The Imm, Mk and Ery1-6 clusters formed a single, branched supercluster when UMAP dimensional reduction was applied (**Fig. 1b**), suggesting that these cell types can share a common hematopoietic progenitor lineage from the immune cell cluster. Instead, hepatocytes and endothelial cells formed segregated clusters from each other and the immune cell cluster, consistent with distinct lineage origins.

### Identifying haematopoietic progenitors in E12.5 liver

We investigated the Imm, Mk and Ery1 clusters in more detail to identify subpopulations of cells that contribute to the immune and erythroid cell population in the E12.5 liver (**Fig. 2a,b**). We identified HSPCs via transcripts for *Cd34* and *Cd93* (coding for AA4.1) (**Fig. 2c; Fig. S2a**)(Yokomizo et al., 2022). DEGs for these cells compared to other liver cells were *Sox4, Gimap1, Hmga2, H2afy, Marcks* (**Fig. 2c; Fig. 3a; Fig. S2a**). Unsupervised clustering divided these HSPCs into 3 separate subclusters, which were annotated based on the expression of additional key marker genes. All three subclusters expressed the HSPC markers *Flt3, Kit* and *Myb* at different levels (**Fig. 2c; Fig. S2a**). By comparing DEGs between these 3 progenitor populations, we identified the cluster with the highest *Ly6a* levels as pre-HSCs/HSCs, the cluster with low *Ly6a* levels but similarly high *Myb* and *Kit* levels as EMP/multipotent progenitor (MPPs) and the cluster with high *Flt3* and *Myb* and *Kit* levels as lympho-myeloid progenitors (LMPs) (**Fig. 2c; Fig. S2a**).

**Figure 2.**
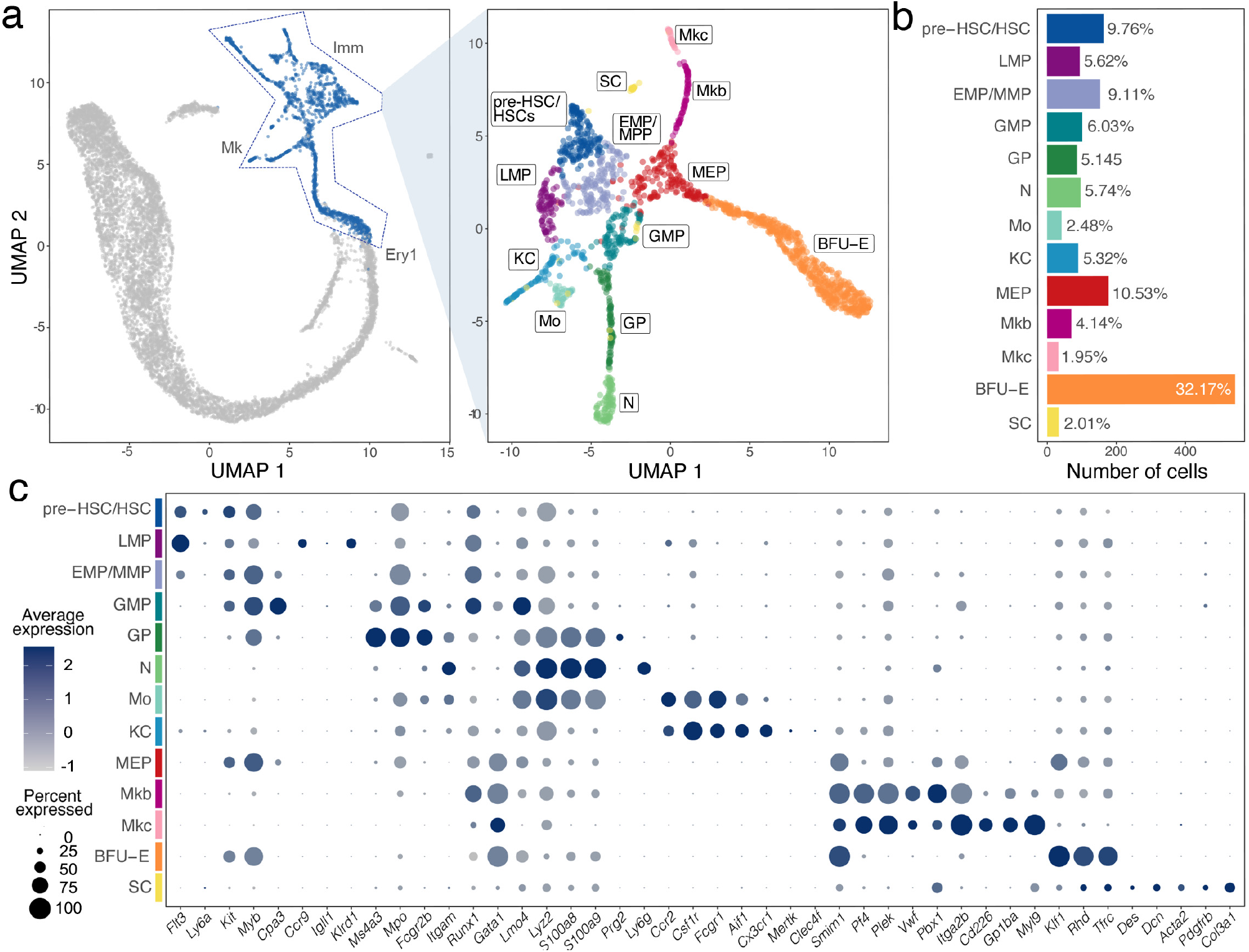
Haematopoietic cell identification in E12.5 mouse liver by scRNA-seq analysis. (**a**) Subset selection followed by UMAP plot visualisation identifies subclusters of distinct hematopoietic cell types. (**b**) A histogram shows the percentage and total number of cells in each subcluster. (**c**) A bubble plot shows expression of selected marker genes in each subcluster; the dot size in represents the percentage of cells within a cell cluster in which a marker was detected, and the dot colour intensity represents the average expression level of that marker.

**Figure 3.**
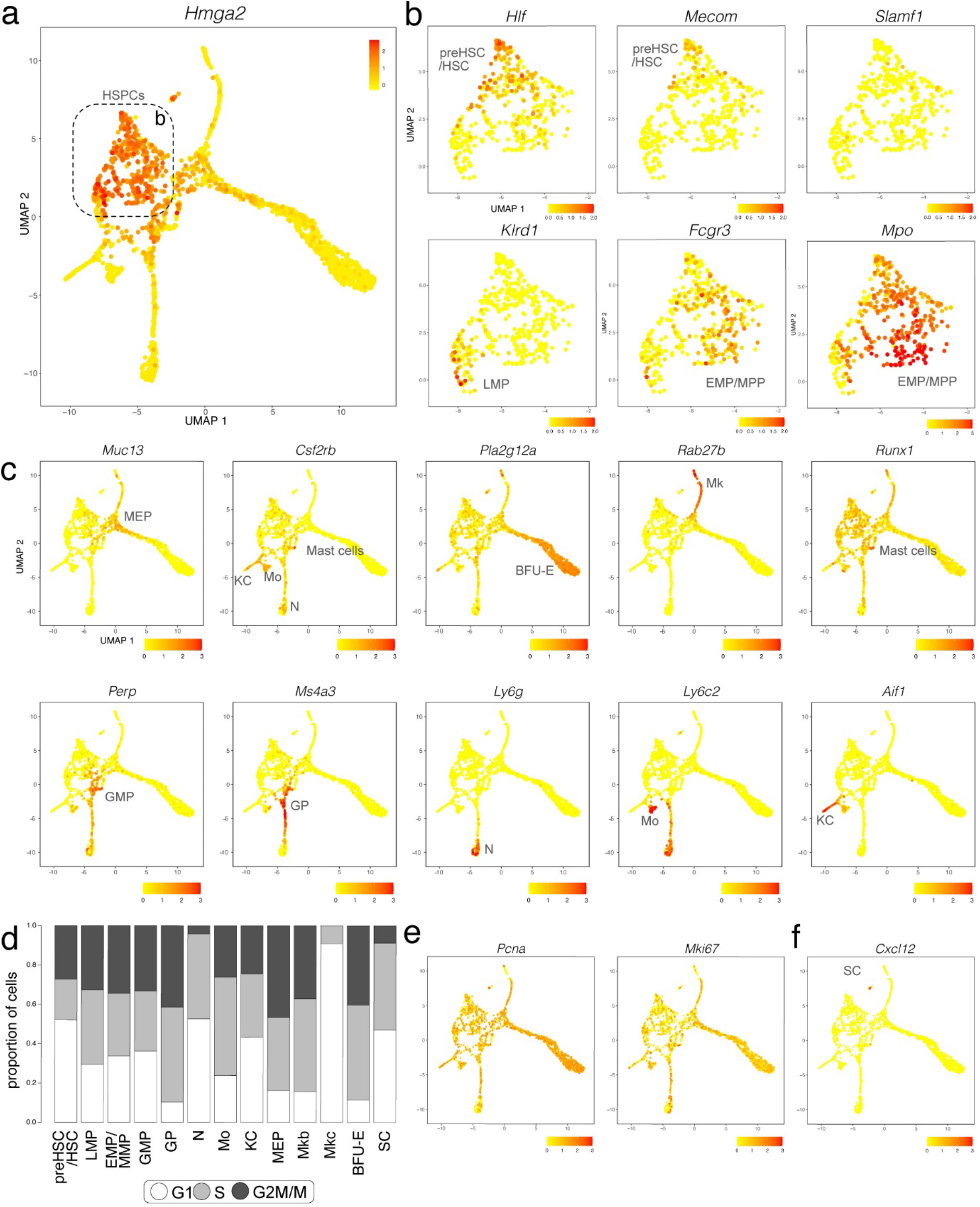
E12.5 mouse liver haematopoietic cells analysed with scRNA-seq data. (**a,b**) A UMAP plot of E12.5 mouse liver haematopoietic cells visualises *Hmga2* expression (**a**), which was used to select haematopoietic progenitor subclusters, indicated with a box and shown at higher magnification in (**b**) for expression of the indicated progenitor genes. Each UMAP plot names the cluster(s) expressing the indicated gene. (**c**) UMAP plots E12.5 mouse liver haematopoietic cells visualise expression of the indicated markers for distinct branches of haematopoietic cell differentiation and hepatic stellate cell progenitors. Each UMAP plot names the cluster(s) expressing the indicated gene. (**d**) The CellCycleScoring prediction algorithm identifies the proportion of cells in G1, S, or G2/M cell cycle phases for each haematopoietic cell cluster. (**e,f**) UMAP plots visualise expression of *Pcna* and *Mki67* (**e**) and *Cxcl12* (**f**).

We further found that the pre-HSC/HSC population had the highest levels of *Hlf*, recently shown to mark the developmental pathway to HSCs but not EMPs (Yokomizo et al., 2019). The pre-HSCs/HSC population was also enriched in *Cd27, Mycn, Hoxa3, Hoxa7, Hoxa9, Rbp1, Hacd4, Cdkn1c* and *Mecom* (**Fig. 3b; Fig. S2a,b**). However, this cell population did not yet express *Slamf1* (encoding CD150) (**Fig. 3b**), a widely accepted marker used to label foetal HSCs at late gestation (Yokomizo et al., 2022). The LMP population included *Ccr9* and *Il7r-positive* cells, as recently shown (Gao et al., 2022) and specifically expressed *Igll1*, *Klrd1* and *Pax5*. The LMP population was also enriched in transcripts for *Ccl3, Ccl4, Pld4, Lsp1, Plac8, Ckb* and *Mndal* but lacked transcripts for the HSC and MPP markers *Hlf* and *Cd48*, respectively (**Fig. 3b; Fig. S2a,b**). The EMP/MMP population was enriched for *Fcgr3* transcripts (encoding CD16) and *Cd48* (**Fig. 3b;S2a,b**), which are known EMP and MPP markers, respectively, but not HSC markers (Soares-da-Silva et al., 2021). The distinctive gene signature of the EMP/MMP population also included *Mpo, Calr, Ccl9, Ap3s1, Cpa3, Anxa3* and *Ctsg* (**Fig. 3b;S2a,b**). The yolk sac EMP marker *Csf1r* (Gomez Perdiguero et al., 2015; Hoeffel et al., 2015) was expressed at low levels in the EMP/MMP population (**Fig. 2c**), but less prominently expressed than in monocytes and Kupffer cells (see below; **Fig. 2c**). By contrast, *Csf1r* transcripts were hardly detected in pre-HSCs/HSCs (**Fig. 2c**). Therefore, *Csf1r* expression in the presence of progenitor markers can be used to distinguish EMPs or other MPPs from pre-HSCs/HSCs.

### Molecular and cellular landscape of megakaryocyte, erythroid and myeloid lineages in E12.5 liver

UMAP dimensionality reduction showed that the EMP/MPP cluster formed a differentiation continuum with two neighbouring cell clusters, the megakaryocyte-erythroid progenitors (MEPs) and granulocyte-monocyte progenitors (GMPs) (**Fig. 2a**). Marker and DEG analysis of all the subclusters within the Imm, Mk and Ery1 populations suggested that MEP and GMP underwent commitment already at E12.5 along the erythro-megakaryocytic and the myeloid lineages, respectively, as described below.

MEPs could be distinguished from other HSPC populations by higher transcript levels for *Myb* and *Kit*, together with the erythroid markers *Gata1* and *Klf1* and the erythroid-megakaryocytic marker *Smim1*. The top DEGs in MEPs were *Muc13* and *Car1*. MEPs formed lineage trajectories towards erythroid progenitors expressing *Klf1, Rhd* and *Tfrc*, identified as BFU-E (see **Fig. 4**), and megakaryoblasts and megakaryocytes expressing *Pf4* and *Plek* (**Fig. 2c; Fig. 3c; Fig. S2a**). The top DEGs in erythroid progenitors were *Pla2g12a, Asns* and *Rexo2*, whereas the top DEGs in megakaryoblasts and megakaryocytes were *F2rl2, Rab27b, Gp1bb, Pbx1, Gp9* and *Rap1b*. Megakaryoblasts were also enriched in transcripts for *F2r, Pbx1, Lat* and *Clec1b*, whereas megakaryocytes were enriched in transcripts for *Myl9, Gp1ba, Itga2b, Cd226, Mest* and *Hist1h2bc* (**Fig. 3c; Fig. S2a**).

**Figure 4.**
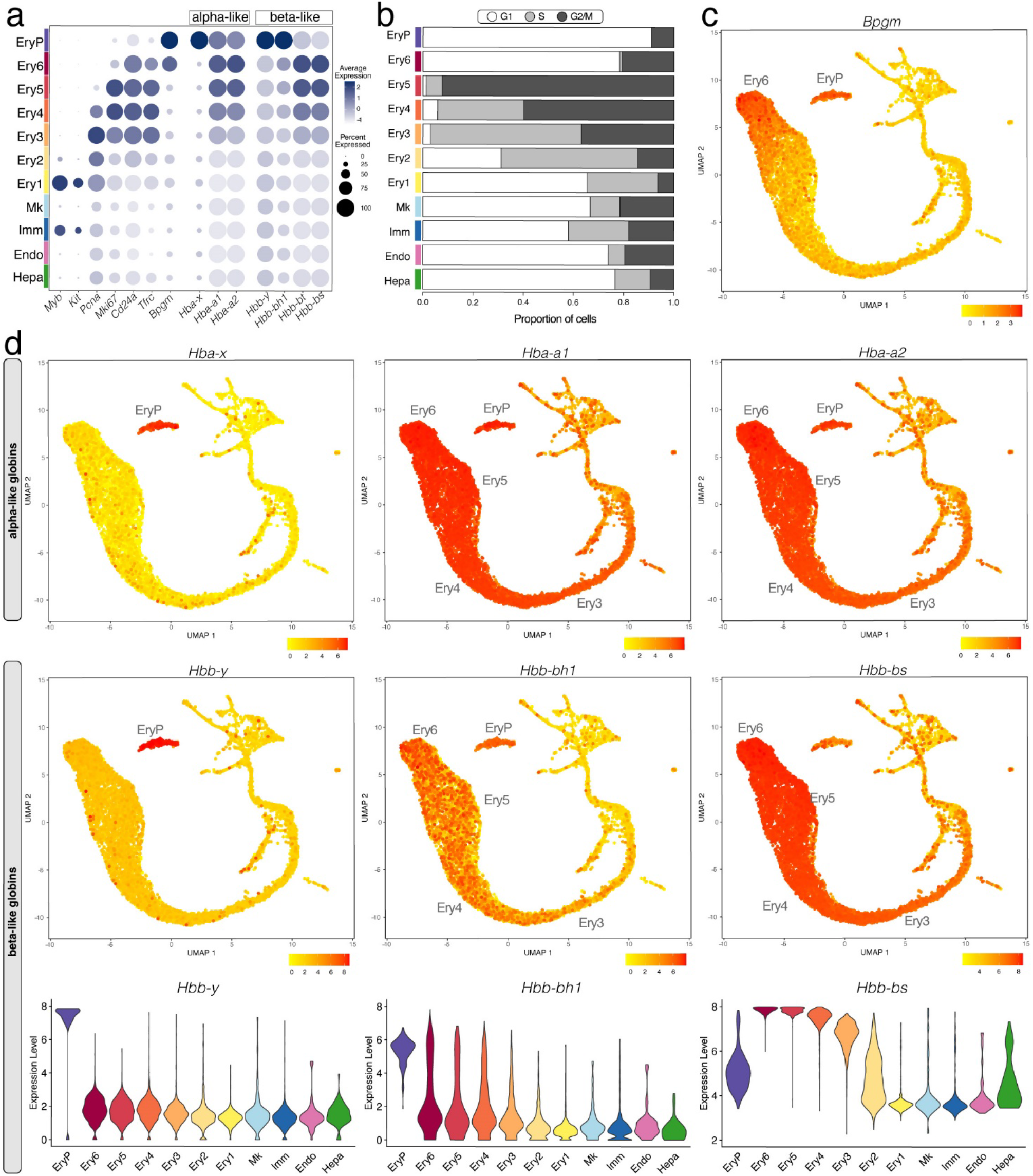
E12.5 mouse liver erythropoiesis analysed with scRNA-seq data. (**a**) A bubble plot shows expression of the selected marker genes in each E12.5 foetal liver cluster; the dot size represents the percentage of cells within a cell cluster in which that marker was detected, and the dot colour intensity represents the average expression level. (**b**) The CellCycleScoring prediction algorithm identifies the proportion of cells in G1, S, or G2/M cell cycle phases for each cluster. (**c**) Expression of the mature erythrocyte marker *Bmpg*. (**d**) UMAP and violin plots visualise the α-like and β-like globin genes expressed in different erythrocyte lineages. In (**c,d**), the cluster(s) expressing the indicated gene are named.

GMPs could be distinguished from other HSPC populations by their enrichment in transcripts for *Hdc, Perp* and *Fcgr3* (**Fig. 3c; Fig. S2a**). The GMP population formed lineage trajectories towards granulocyte progenitors enriched in *Ms4a3, Mpo* and *Fcgr2b*, with the top DEGs being *Cd63* and *Elane* (**Fig. 2c; Fig. 3c; Fig. S2a**). Granulocyte progenitors then led to neutrophils/granulocytes, which were enriched in transcripts for *Itgam* encoding CD11b, *S100a8, S100a9* and *Ly6g;* other top DEGs were *Retnlg, Camp, Ngp, Ltf, Lcn2, Ifitm6* and *Stfa1* (**Fig. 2c; Fig. 3c; Fig. S2a**). A parallel lineage trajectory led from GMPs to Kupffer cells/tissue macrophages, which were enriched in transcripts for *Aif1* encoding IBA1, *Cx3cr1, Mertk* and *Clec4f*, and to monocytes, which were enriched in transcripts for *Ccr2, Ly6c2 and Pld4* (**Fig. 2c; Fig. 3c; Fig. S2a**). Monocytes and granulocyte progenitors shared high levels of *Lgals3* and *Fcnb* transcripts with each other and further shared *Lyz2* and *Hp* transcripts with neutrophils (**Fig. 2c; Fig. S2a**). *Csf1r* transcripts were high in Kupffer cells/tissue macrophages and even higher in monocytes, but not detected in differentiated neutrophils (**Fig. 2c**). *Csf1r* expression can therefore be used as a distinguishing marker for the macrophage/monocytic branch within the myeloid lineage. Cells enriched in mast cell markers *Cpa3* and *Runx1* transcripts (Grootens et al., 2018) appeared to bud from the GMP cluster (**Fig. 2e**) and may represent early foetal mast cells.

### Cell cycle analysis of haematopoietic lineages in the E12.5 liver

The CellCycleScoring prediction algorithm in Seurat together with expression analysis for the S marker *Pcna* (Kurki et al., 1986) and S/G2 marker *Mki67* (Miller et al., 2018) reported that cell number expansion mainly occurs in MEPs as well as downstream of MEPs in the megakaryoblasts and BFU-E of the megakaryocyte/erythroid branch as well as in granulocyte progenitors of the myeloid branch (**Fig. 3d,e**). Cell number expansion was more moderate in HSPCs at E12.5, with pre-HSCs/HSCs appearing less proliferative than other progenitors at this stage (**Fig. 3d,e**). Liver megakaryocytes appeared to have already terminally differentiated at E12.5, because this cluster lacked cells entering the G2/M phase, with less than 10% of cells in this cluster in S phase (**Fig. 3d,e**). Within the myeloid lineage, neutrophils also appeared already terminally differentiated, whereas the majority of Kupffer cells and especially monocytes were either in the S or G2/M phase (**Fig. 3d,e**).

### Molecular and cellular landscape of erythropoiesis in E12.5 liver

Mouse foetal liver erythroid cells were identified by their expression of haemoglobin genes, the erythroid-specific transcription factor *Klf1* and the erythroid membrane marker *Rhd* (**Fig. 1d**). We first examined the expression pattern of the nuclear long noncoding RNA (lncRNA) *Malat1* as a nucleus marker (Yu and Shan, 2016). As *Malat1* was present in all cells of the erythroid clusters identified in the E12.5 liver, they all contained a nucleus (**Fig. S1d**), consistent with an origin from yolk sac-born primitive erythrocytes circulating through the liver (EryP cluster) and from liver-resident transient-definitive erythroid progenitors and erythroblasts (Ery1-6 clusters). The total number of expressed genes decreased gradually from Ery3 to Ery5, resulting in a drastic reduction of transcriptomic complexity in both Ery6 and EryP (**Fig. S1e**), agreeing with an erythroid maturation program that includes enhanced chromatin condensation and consequent transcription silencing (Chen et al., 2009).

CellCycleScoring prediction together with *Pcna* and *Mki67* expression analysis suggested that no cells in the EryP cluster were in S phase and fewer than 10% in G2/M (**Fig. 4a,b**), consistent with them being terminally differentiated erythrocytes. The Ery2-5 clusters, instead, contained the most proliferative cells in the whole liver (**Fig. 4a,b**), suggesting rapid expansion of erythroid progenitors or erythroblasts in the E12.5 liver. Cells in the Ery6 cluster were mostly in G1 phase similar to EryP cluster, with no cells in S phase and 20% in G2/M (**Fig. 4a,b**) and therefore likely represent orthochromatic erythroblasts that had ceased to divide (Hasan et al., 2018)

Gene expression analysis showed that the Ery1 cell population contained transcripts for *Myb* and *Kit* but not *Cd24a*, and thus likely corresponds to BFU-E progenitors. The Ery2 cell population expressed higher levels of *Cd24a* compared to Ery1, but remained haemoglobin-negative, and thus likely corresponds to CFU-E progenitors. Transcripts for *Cd24a, Tfrc* encoding CD71 and haemoglobin genes increased from the Ery3 to the Ery4 and then Ery5 cell populations (**Fig. 4a**), similarly to what recently reported by flow cytometry (Soares-da-Silva et al., 2021). These populations therefore likely correspond to sequential erythroid stages proerythroblasts, basophilic erythroblasts and polychromatophilic erythroblasts. Orthochromatic erythroblasts (Ery6 cluster) and primitive erythrocytes (EryP) both expressed the mature erythrocyte marker *Bpgm*, but orthochromatic erythroblasts contained transcripts for *Cd24a* and *Tfrc*, whereas these were barely detectable in primitive erythrocytes (**Fig. 4a,c**). *Bpgm* together with *Cd24a* and *Tfrc* expression could therefore be used as distinguishing markers between orthochromatic erythroblasts and primitive erythrocytes.

### Haemoglobin gene expression profiles in the E12.5 liver

In the E12.5 liver, transcripts for the embryonic α-like globin *Hba-x* (ζ) and β-like globin *Hbb-y* (εy) were restricted to the EryP cluster, which also expressed the α globins *Hba-a1* and *Hba-a2* (α) as well as the β-like globin *Hbb-bh1* (βH1), but low levels of adult β globins *Hbb-bs* and *Hbb-bt* (**Fig. 4a,d**; note that *Hbb-bs* and *Hbb-bt* are specifically present in C57BL/6 mice, whereas BALB/c and 129Sv mice have a different haplotype that includes *Hbb-b1* and *Hbb-b2*). To corroborate that the EryP cluster represents primitive erythrocytes, we also analysed scRNA-seq data from whole embryos with yolk sacs at E8.5 (Pijuan-Sala et al., 2019), when primitive erythrocytes arise in the yolk sac (Palis et al., 1999). EryP at E8.5 were identified by *Klf1* and *Rhd* (**Fig. S3a,b**) and were nucleated (*Malat1*-positive; **Fig. S3c**). Similar to E12.5, EryP at E8.5 had high levels of *Hba-x* and the *Hba-a1* and *Hba-a2* (α) as well as *Hbb-y* and *Hbb-bh1* (βH1), but barely detectable levels of adult β globin *Hbb-bs* and *Hbb-bt* (**Fig. S3d**). By contrast to EryP cells at E12.5, however, EryP at E8.5 still expressed low transcript levels for the mature erythrocyte marker *Bpgm* (**Fig. S3d**). The adult β globin transcripts *Hbb-bs* and *Hbb-bt* were highly expressed in the erythroid clusters Ery3, Ery4, Ery5, Ery6 of E12.5 liver alongside the α globins *Hba-a1* and *Hba-a2*, (**Fig. 4a,d**). These clusters also retained *Hbb-bh1* expression, although at lower levels than in primitive erythrocytes (EryP) (**Fig. 4a,d**). Foetal liver erythropoiesis can therefore be distinguished from primitive erythrocytes by the expression of both adult β globin genes and the embryonic β-like globin *Hbb-bh1*.

### Onset of foetal liver haematopoiesis

To better understand the onset of haematopoiesis in the mouse foetal liver, we performed a time-course single cell analysis with published scRNA-seq dataset of mouse foetal liver cells at E9.5, E10.5 and E11.5 (Dong et al., 2018).

In the E9.5 mouse liver, we identified *Afp*-positive hepatocytes and *Col3a1*-positive mesenchymal cells, but *Klf1*-positive erythroid and *Cx3cr1*-positive myeloid cells could not be detected at this stage, with just one cell out of the total 92 cells expressing the EMP markers *Kit*, *Itga2b, Cd34* and *Csf1r* (**Fig. S4a**, arrowhead). At E10.5, we identified many haematopoietic progenitors expressing the EMP markers *Cd34*, *Csf1r*, *Mpo*, *Fcgr3* and *Ptprc*, and they formed a lineage continuum with myeloid cells expressing *Cx3cr1* and *Csf1r* and megakaryoblasts expressing *Itga2b* (**Fig. S4b,c**). Haematopoietic progenitors (including EMPs) began to express transcripts for *Myb, Fcgr2b* (coding for CD32) and the MPP marker *Cd48* by E11.5 while downregulating transcripts for *Kit* and *Itga2b* (**Fig. S4b,c**).

Coincident with liver homing by EMPs from E10.5 onwards, *Klf1*-expressing erythroid clusters could be detected in the foetal liver from E10.5 onwards and increasingly at E11.5, when they were clearly distinguishable from *Hba-x* positive primitive erythrocytes (EryP) (**Fig. S4b,c**). The liver-derived erythroid cells at both E10.5 and E11.5 included *Myb-* and *Kit*-positive cells (Ery1; **Fig. S4b,c**), as well as *Cd24a*-expressing cells that did not express any haemoglobin genes (Ery2; **Fig. S4b,c**), indicating that they are BFU-E and CFU-E, respectively. A separate cluster of erythroid cells expressing *Tfrc* and low levels of *Hbb-bh1*, representing the first foetal erythroblasts, could be identified in the E11.5 liver (Ery3; **Fig. S4c**), further validating the existence of a foetal intermediate haemoglobin profile.

We validated our observations by analysing another publicly available dataset that includes 14597, 27998 and 16592 foetal liver cells at E11.0, E11.5 and E13.0, respectively (Wang et al., 2020). At all three stages, haematopoietic progenitors expressing *Kit*, *Myb*, *Cd34, Mpo*, and *Ptprc* formed a lineage continuum with MEPs expressing *Muc13*. At both E11.0 and E11.5, liver MEPs expressed *Csf2rb*, a marker for yolk sac MEPs destined to colonise the foetal liver (Iturri et al., 2021). At these stages, *Csf2rb* expression was also observed in HSPCs (except LMPs) as well as in myeloid cells (**Figs. 5a-c** and **6a-c**). At E12.5 and E13.0, *Csf2rb* transcripts remained abundant in myeloid lineages and, especially, in mast cells, but had been lost from liver MEPs (**Figs. 3c** and **7a-c**). MEPs further branched into *F2r*-positive megakaryoblasts followed by *Myl9*-expressing megakaryocytes (**Fig. 5a-c; Fig. 6a-c; Fig. 7a-c**) and *Klf1-* and *Myb*-positive but *Cd24a*-negative BFU-E erythroid progenitors, which led via *Cd24a*-positive CFU-E progenitors to the first erythroblasts with *Hbb-bh1* and *Hbb-bs* haemoglobin expression (**Fig. 5a-d; Fig. 6a-d; Fig. 7a-d**).

**Figure 5.**
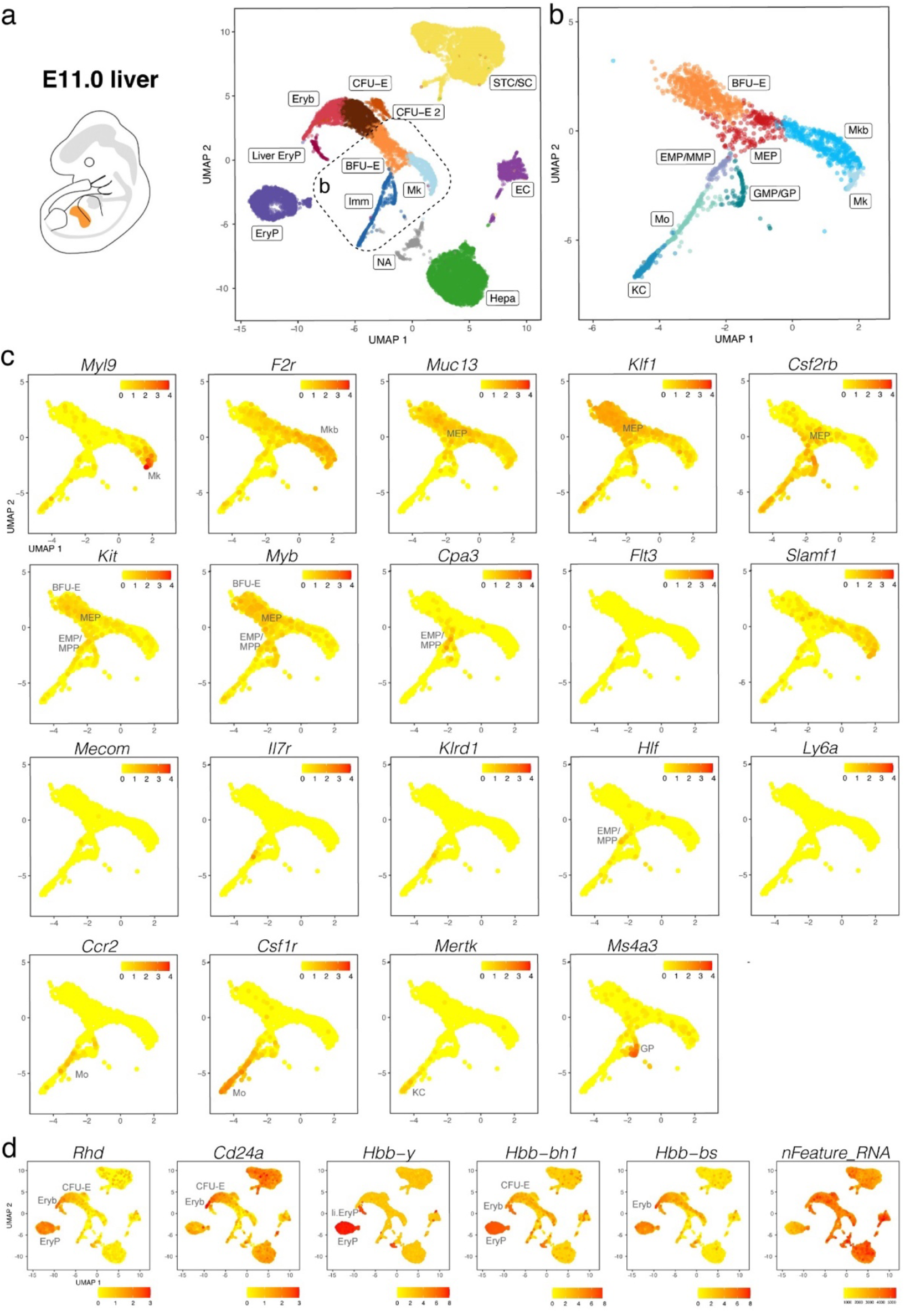
scRNA-seq of the E11.0 mouse liver. (**a,b**) The UMAP plot in (**a**) visualises clusters of distinct cell types in the total dataset; the box indicates the Mk, BFU-E and Imm clusters, which were subclustered in (**b**). (**c,d**) UMAP plots visualise expression of the indicated genes that serve as markers of distinct branches of haematopoietic cell (**c**) and erythroid differentiation (**d**). In (**c,d**), each UMAP plot names the cluster(s) expressing the indicated gene. EC, endothelial cells; li. EryP, liver primitive erythrocytes; Mkb, megakaryoblasts; NA, not assigned; STC/SC, septum transversum cells/hepatic stellate cells.

**Figure 6.**
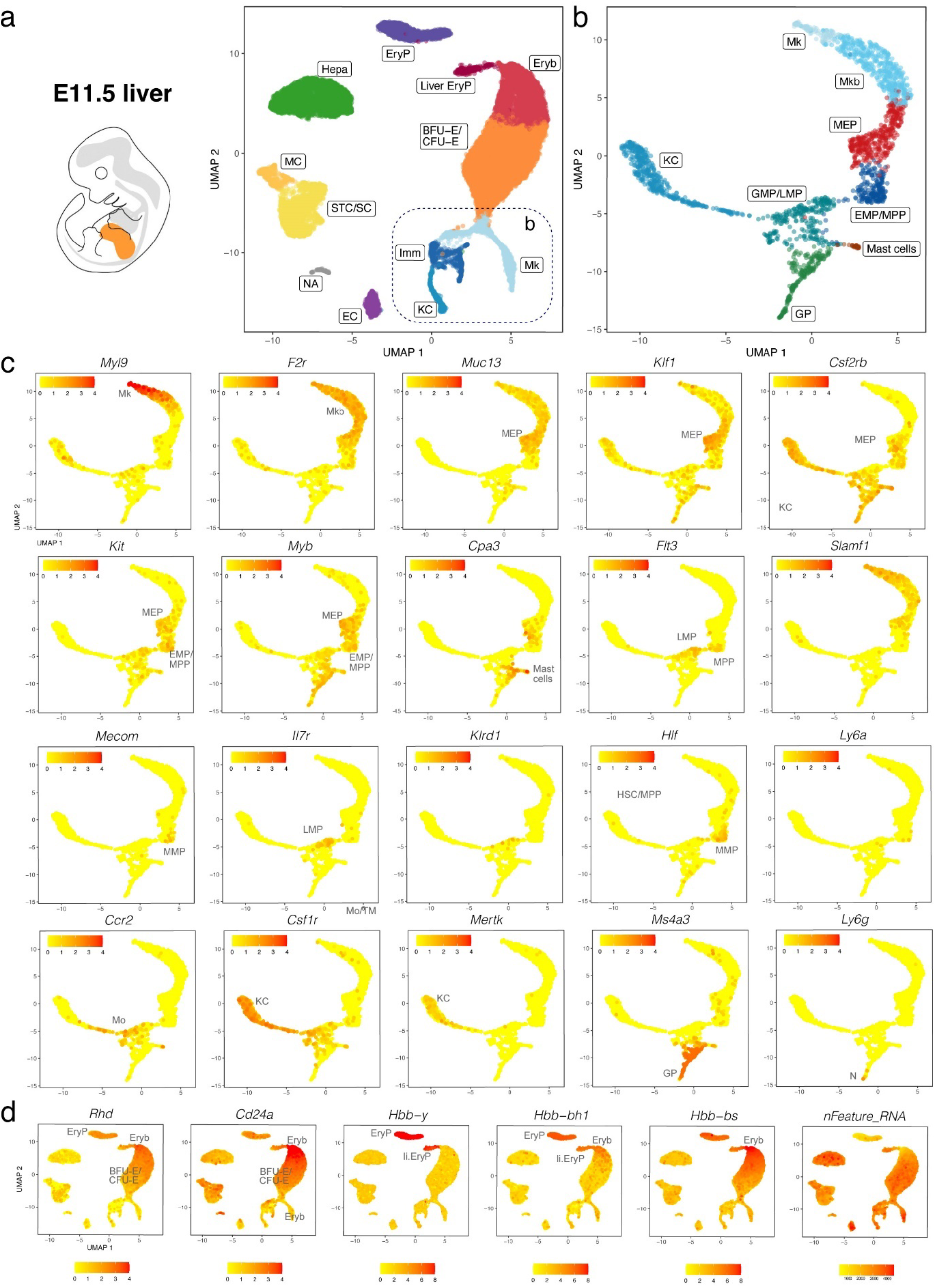
scRNA-seq of the E11.5 mouse liver. (**a,b**) The UMAP plot in (**a**) visualises clusters of distinct cell types in the total dataset; the box indicates the Mk, KC and Imm clusters, which were subclustered in (**b**). (**c,d**) UMAP plots visualise expression of the indicated genes that serve as markers of distinct branches of haematopoietic cell (**c**) and erythroid differentiation (**d**). In (**c,d**),each UMAP plot names the cluster(s) expressing the indicated gene; MC, mesothelial cells.

**Figure 7.**
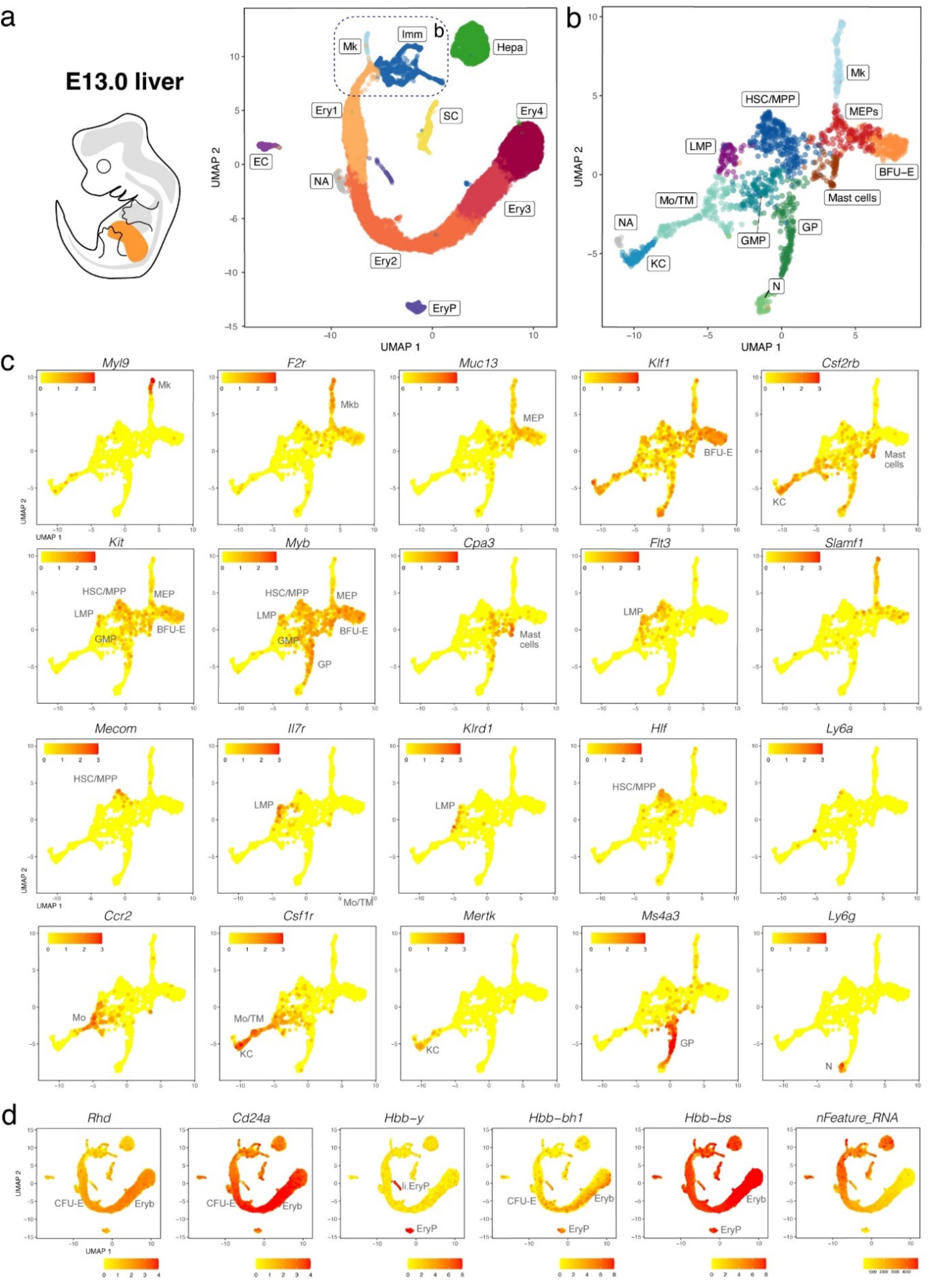
(**a,b**) The UMAP plot in (**a**) visualises clusters of distinct cell types in the total dataset; the box indicates the Mk, BFU-E and Imm clusters, which were subclustered in (**b**). (**c,d**) UMAP plots visualise expression of the indicated genes that serve as markers of distinct branches of haematopoietic cell (**c**) and erythroid differentiation (**d**). In (**c,d**), each UMAP plot names the cluster(s) expressing the indicated gene; SC, hepatic stellate cells.

At all three stages, we identified primitive erythroid cells with reduced transcriptomic complexity that were enriched in embryonic but not adult globins and likely are yolk sac-born primitive erythrocytes circulating through the liver (EryP cluster, **Fig. 5d; Fig. 6d; Fig. 7d**; compare to **Fig. 4; Fig. S1e**). Additionally, at E11.0 and E11.5, a small population of erythroid cells expressed a higher number of genes, and they were enriched in embryonic but not adult haemoglobin genes; further, their cluster proximity in the UMAP suggested that their transcriptomic signature is more similar to that of liver-derived erythroblasts than yolk sac-derived primitive erythrocytes (liver EryP; **Fig. 5d; Fig. 6d**). These erythroid cells could not be identified in our E12.5 dataset (**Fig. 4c,d**), and very few were detected in the E13.0 dataset (**Fig. 7d**), suggesting that they might be the product of short-lived primitive erythroid progenitors that had homed to the foetal liver before E11.0.

Haematopoietic progenitors also formed a lineage continuum with myeloid cells, including monocytes expressing *Ccr2*, Kupffer cells expressing *Mertk* and granulocyte progenitors expressing *Ms4a3* (**Fig. 5a-c; Fig. 6a-c; Fig. 7a-c**). Granulocyte progenitors were already present at E11.0, but the first *Ly6g*-expressing neutrophils/granulocytes appeared only at E11.5 (**Fig. 5b,c; 6b,c**). From E11.5 onwards, mast cells formed a cluster distinct from GMPs, and at E13.0, the mast cell cluster had become larger and appeared phenotypically closer to MEPs when compared to earlier stages (**Fig. 5b,c; Fig. 6b,c; Fig. 7b,c**). We identified two distinct subsets of tissue macrophages in the E13.0 dataset. One cluster with high *Mertk* and *Csf1r* levels was already present at earlier stages and represented Kupffer cells (KC; **Fig. 7a-c**), which are of yolk sac origin (Ginhoux and Guilliams, 2016). A second subset of tissue macrophages expressing *Csf1r*, *Adgre1* and *Cx3cr1*, but not the monocyte marker *Ccr2* or the mature KC marker *Mertk*, was present within a cluster that also contained *Ccr2*-positive but *Adgre1*-negative monocytes (Mo/TM; **Fig. 7a-c**). This subset likely represents liver monocyte-derived tissue macrophages that will gradually replace the initial pool of yolk sac-derived Kupffer cells during late gestation (Ginhoux and Guilliams, 2016).

*Ly6a*-positive HSCs or pre-HSCs were rare in E12.5 or E13.0 mouse foetal liver (**Fig. 2c; Fig. 7c**), and could not be identified at earlier stages (**Fig. 5c; Fig. 6c**). Further, HSCs expressing *Slamf1* (encoding CD150) were not detected at any time point up to E13.0 (**Fig. 5c; Fig. 6c; Fig. 7c**). By contrast, rare *Flt3-, Hlf*- and *Mecom*-positive MPPs as well as *Il7r-* and *Klrd1*-positive LMPs could be identified at all stages (**Fig. 5c; Fig. 6c; Fig. 7c**). Prior to E12.5, foetal liver haematopoiesis is therefore predominantly driven by transient definitive progenitors first appearing in the mouse liver at E10.5, rather than definitive progenitors.

### Non-haematopoietic cells in the E12.5 liver

In addition to haematopoietic cells, the foetal liver contains structural cell types crucial for liver growth and function. These cells segregated apart from the blood and immune cells in our scRNA-seq dataset of the E12.5 liver into 3 main clusters.

One main cell cluster had high levels of hepatoblast markers such as *Afp* and *Alb*, but lacked the mature hepatocyte marker *Epcam* (Hepa cluster; **Fig. 1b-d**). Unsupervised subclustering identified two separated subclusters (**Fig. S5a**). One subcluster had higher levels of hepatoblast markers (*Afp*, *Alb, Hnf4a, Krt8*) than the other and also upregulated the hepatocyte differentiation markers (*Tbx3, Cebpa* and *Prox1;* **Fig. S5b**). Cells in this cluster are likely hepatoblasts undergoing hepatocyte differentiation. The other subcluster had reduced expression of hepatoblast markers (*Afp*, *Alb*) and lacked transcripts for other hepatoblast markers (*Hnf4a, Krt8*) but expressed *Spp1*, a marker for cholangiocytes (Yang et al., 2017) (**Fig. S5b**). These cells are likely hepatoblasts undergoing cholangiocyte differentiation. Analysis of differentially expressed genes (DEGs) showed that the presumed cholangiocyte precursors were enriched in *Cct4, Cct7, Eif4a1, Ptma* and *Actb*, whereas *Gpc3, Hpx, Apoh, Serpina1c, Rrbp1, Igf2, Fgg, Meg3* and *Elovl2* were underrepresented when compared to hepatoblasts (**Fig. S5c**).

Hepatic stellate cell precursors enriched in *Pdgfrb, Acta2, Dcn, Cxcl12, Col3a1* and *Des* (**Fig. 2a-c; Fig. 3f**) formed another separate cluster with other DEGs being *Nnat, Col1a2, Ptn, Sparc, Meg3* and *H19* (**Fig.S2a**).

A third cell cluster, the Endo cluster, was representative of liver sinusoidal endothelial cells (LSECs; **Fig. 1b-d**), because transcripts for the endothelial markers *Cldn5* and *Sox18* were present together with core LSEC genes such as *Lyve1* and *Plvap*. Unsupervised subclustering revealed the presence of 2 distinct cell populations, namely LSEC1 and LSEC2 (**Fig. S5a**). Both subclusters lacked transcripts for the periportal LSEC markers *Tm4sf1* and *Clec14a* (**Fig. S5d**), previously identified in adult human liver (MacParland et al., 2018). The LSEC1 subcluster appeared enriched for *Oit3* and *Cldn5*, recently reported to be increased in LSECs from central vein proximity (MacParland et al., 2018). They were also enriched for *Eng*, which is upregulated in endothelial cells from large calibre vessels in the adult liver, such as the central veins (MacParland et al., 2018). Further, the LSEC1 subcluster was enriched for *Ephb4*, an Eph receptor selectively expressed on vein endothelial cells (Gerety et al., 1999) (**Fig. S5e**). Other DEGs for LSEC1 include *Asb4, Ppp1r14b, Cnbp, Col4a2, Plk2, Polr2e, Tubb6, Plpp3, Lpar6*) (**Fig. S5c**). By contrast, the top DEG for LSEC2 was *Bex2* (**Fig. S5c,e**). These findings suggest that at E12.5 LSECs were composed by endothelial cells that either began acquiring a central vein zonation phenotype (LSEC1) or lacked a zonation-specific phenotype (LSEC2).

## Discussion

Defining the mouse embryonic liver environment at single-cell resolution using scRNA-seq will inform whether and how the mouse provides a suitable model to understand the molecular bases of human liver development, function and disease, including congenital immunodeficiencies, anaemia and also childhood leukaemia (Cazzola et al., 2020). Here we addressed limitations in prior studies that either performed a preselection step to enrich the dataset only for specific cell types (Freyer et al., 2020; Gao et al., 2022) or included only a small number of all foetal liver cell types (Dong et al., 2018; Su et al., 2017) and extended the analysis of a study that obtained the transcriptomes of a larger number of liver cells but analysed mostly hepatocyte development (Wang et al., 2020).

Consistent with current knowledge generated by lineage tracing in combination with histology and flow cytometry (Barone et al., 2022; Canu and Ruhrberg, 2021), we found that haematopoietic cells were rare in the foetal liver at E9.5 but consistently present in this organ from E10.5 onwards (**Fig. S4**). Further, our data agree with erythroid cells derived from liver-resident, transient-definitive progenitors becoming the main cellular component of the foetal liver by E12.5 (**Fig. 1**). We observed that transient-definitive erythroblasts generated in the foetal liver retained expression of the embryonic β-like globin *Hbb-bh1* (βH1) together with α globins *Hba-a1* and *Hba-a2*, although at lower levels than in primitive erythrocytes (**Fig. 4**). This is interesting, because it was previously suggested that intermediate foetal haemoglobin composed by α globins and a foetal β-like globin is a specific feature of anthropoid primates, with βH1 globin expressed only by primitive erythrocytes in the mouse (Dzierzak and Philipsen, 2013). Our observations instead agree with recent descriptions of low βH1 transcript levels in the mouse foetal liver (McGrath et al., 2011; Soares-da-Silva et al., 2021). These prior studies used *in situ* hybridisation and flow cytometry analyses to detect βH1 transcripts, but these techniques were not combined with markers suitable to clearly distinguish liver transient-definitive erythroblasts from primitive erythrocytes that circulate through the foetal liver, where both types of erythroid cells become enucleated following interaction with macrophages in the erythroblastic islands. These limitations are readily addressed by scRNA-seq data, because the two types of erythroid cells segregated in two distinct clusters (**Figs. 4,6,7**). Our scRNA-seq analysis also identified a short-lived population of erythroid cells at in E11.0 and 11.5 liver; these were phenotypically similar to liver-derived erythroblasts and clustered separately from primitive erythrocytes, despite being enriched in primitive but not definitive haemoglobin genes (**Figs. 5,6**). These cells might be a hitherto unidentified progeny of a late primitive erythroid progenitor or, alternatively, of an early EMP-like progenitor that colonises the early foetal liver but are rapidly replaced when transient definitive erythropoiesis takes hold.

Erythroid, megakaryocytic and myeloid cells each emerged in a separate, continuous trajectory from the HSPC cluster in all datasets analysed between E10.5 and E13.0 (**Figs. 1,2, and 4–7, Fig. S3**). This observation suggests that these cell types share a common hematopoietic progenitor in the foetal liver up to E13.0. HSPCs shared expression of *Cd34, Cd93* (coding for AA4.1), *Hmga2, Flt3, Kit* and *Myb*. By contrast, up to E13.0, cells within the HSPC cluster did not contain transcripts for *Slamf1*, encoding CD150 (**Fig. 2**), which is a widely accepted marker for foetal HSCs. Notably, the *Hlf-* and *Ly6a*-expressing preHSC/HSC subset within the HSPC cluster did not appear more proliferative than other progenitors at E12.5 (**Fig. 3**). As HSCs expand in the foetal liver but minimally contribute to foetal haematopoiesis (Yokomizo et al., 2022), our observations indicate that the E12.5 liver is mostly populated by pre-HSCs or short-term HSCs (ST-HSCs) rather than definitive HSCs, as previously suggested (Rybtsov et al., 2011).

Cell cycle analysis further suggested that haematopoietic cell expansion in the early foetal liver mainly occurs downstream of HSPCs at the level of bipotent or monopotent progenitors, such as MEPs, megakaryoblasts, BFU-E progenitors and granulocyte progenitors (**Fig. 3**). Cell expansion appeared to occur similarly in MEPs as well as megakaryoblasts and BFU-E progenitors in the megakaryocyte/erythroid branch (**Fig. 3**). Instead, different branches of the myeloid lineage showed divergent expansion strategies. Thus, granulocyte progenitors actively proliferated to produce mature neutrophils/granulocytes that had exited the cell cycle (**Fig. 3**). This observation may indicate that these cells expand in the foetal liver before being released into the blood stream. By contrast, differentiated Kupffer cells and monocytes were highly proliferative (**Fig. 3**). It is conceivable that Kupffer cells expand concomitantly with rapid liver expansion to maintain sufficient density, whereas monocytes proliferate to provide enough circulating progenitors for tissue-resident macrophages in peripheral tissues, such as the lung, heart and skin.

Separate from the haematopoietic cells, the progenitors of hepatocytes and also endothelial cells each formed segregated, small clusters. The small cluster size was likely explained by cell types being underrepresented following cell isolation, due to them being bound into cell sheets that need dissociating and are thus subject to loss and damage, compared to naturally singular haematopoietic cells (**Fig. 1**). It was previously reported that hepatoblasts start differentiating into hepatocytes and cholangiocytes at around E13.5 in mouse, with cholangiocyte precursors being generated at the ductal plate from a monolayer of hepatoblasts surrounding the portal veins, whereas hepatoblasts located away from portal vein areas differentiate into hepatocytes later on (Gordillo et al., 2015). Our observations instead suggest that the specification of cholangiocyte precursors from the common hepatoblast progenitor already starts at E12.5. The endothelial cell cluster identified in our scRNA-seq of whole E12.5 mouse foetal liver revealed 2 distinct cell populations, namely LSEC1 and LSEC2 (**Fig. S5**). LSEC1 appeared polarised towards a central vein but not periportal fate, likely reflecting the remodelling pattern already occurring in the liver bud. LSEC2 instead lacked a zonation-specific phenotype. An absent periportal LSEC phenotype might be explained by incomplete remodelling of the right and left vitelline veins into the portal vein or the need for a switch from foetal to postnatal circulation.

Other non-haematopoietic cells in the E12.5 foetal liver are mesenchymal in nature, such as the precursors of hepatic stellate cells (**Fig. 2**). Hepatic stellate cells originate from the septum transversum-derived mesothelium during development and localise in the space of Disse in the adult liver, where they constitute the major mesenchymal component to support liver homeostasis; when activated by injury or infection, stellate cells are the major cell type responsible for liver fibrosis. We found that foetal liver stellate cells were highly enriched in *Cxcl12* transcripts, consistent with their role in recruiting CXCR4-expressing haematopoietic progenitors (Gordillo et al., 2015).

Owing to its genetic tractability, the mouse is the most widely used mammalian model system to understand human development, including liver morphogenesis and haematopoiesis. Although these developmental processes are generally considered conserved across vertebrates (Gordillo et al., 2015; Jagannathan-Bogdan and Zon, 2013), there are notable differences between mouse and human, for example in the expression and function of the surface molecules used to immunophenotype haematopoietic progenitors and mature immune cells (Ivanovs et al., 2017; Parekh and Crooks, 2013). Accordingly, defining haematopoietic development in the mouse foetal liver at single cell level provides an essential resource for better understanding the developmental dynamics that might be affected in congenital or paediatric immune diseases.

## Availability of data and material

The E12.5 foetal liver scRNA-seq has been deposited in NCBI’s Gene Expression Omnibus (NCBI-GEO, GSE180050). Other publicly available datasets analysed in this study include: whole mouse embryo scRNA-seq at E8.5 (ArrayExpress, E-MTAB-6967); foetal liver scRNA-seq at E9.5, 10.5 and 11.5 (NCBI-GEO, GSE87038); foetal liver scRNA-seq at E11.0, 11.5 and 13.0 (CNCB-NGDC, CRA002445).

## Code availability

Not applicable

## Authors’ contributions

SM, CR and AF conceived and designed the study. FS and AF performed mouse experiments. EC, EV and AF performed bioinformatic analyses. CR and AF co-wrote the manuscript. All authors read and approved the submitted manuscript.

## Acknowledgements

We thank Luca Rotta and IEO Genomics Unit, Simona Ronzoni and IEO Flow Cytometry Unit, Elisabetta Dejana and Monica Corada for help with sample processing and sequencing. We thank Giovanni Canu and Emanuele Azzoni for helpful comments on the manuscript. Figure 1a was created with a licence from BioRender.

