## Supplemental Figures for "A refined single cell landscape of haematopoiesis in the mouse foetal liver"

### Supplemental Materials

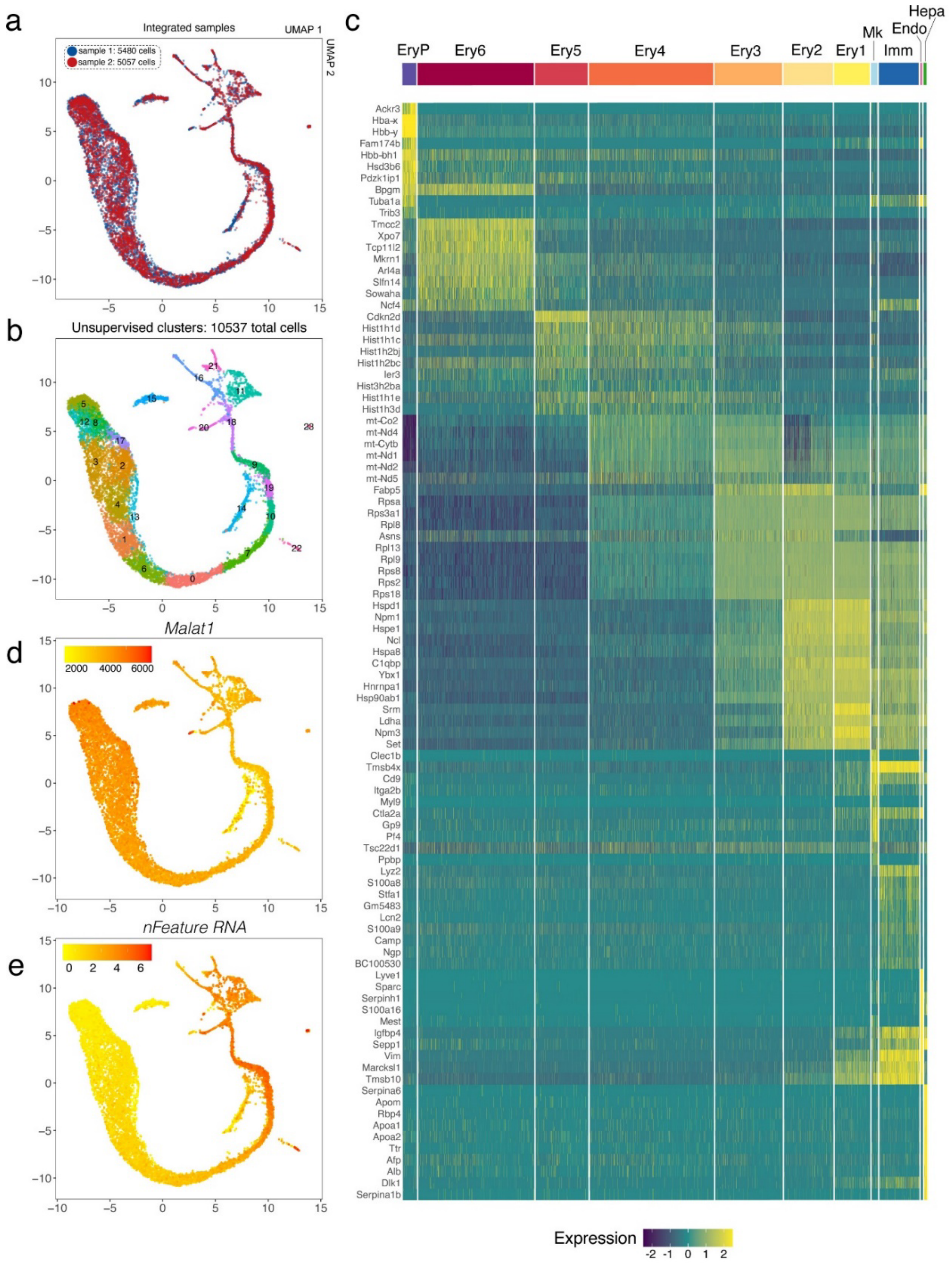


**Figure S1**. **Characterisation of the E12.5 mouse liver scRNA-seq dataset.**
(**a**) A UMAP plot visualises the distributions of the two technical replicates from the same liver sample. (**b**) Unsupervised clustering of cell types into clusters 1-23. (**c**) A heatmap plot of differentially expressed genes (DEGs) among the curated cell clusters. (**d**) A UMAP plot visualises *Malat1*expression; *Malat1*-negative cells in the Ery2 cluster likely represent apoptotic bodies. (**e**) A UMAP plot visualises the number of expressed genes per cell.

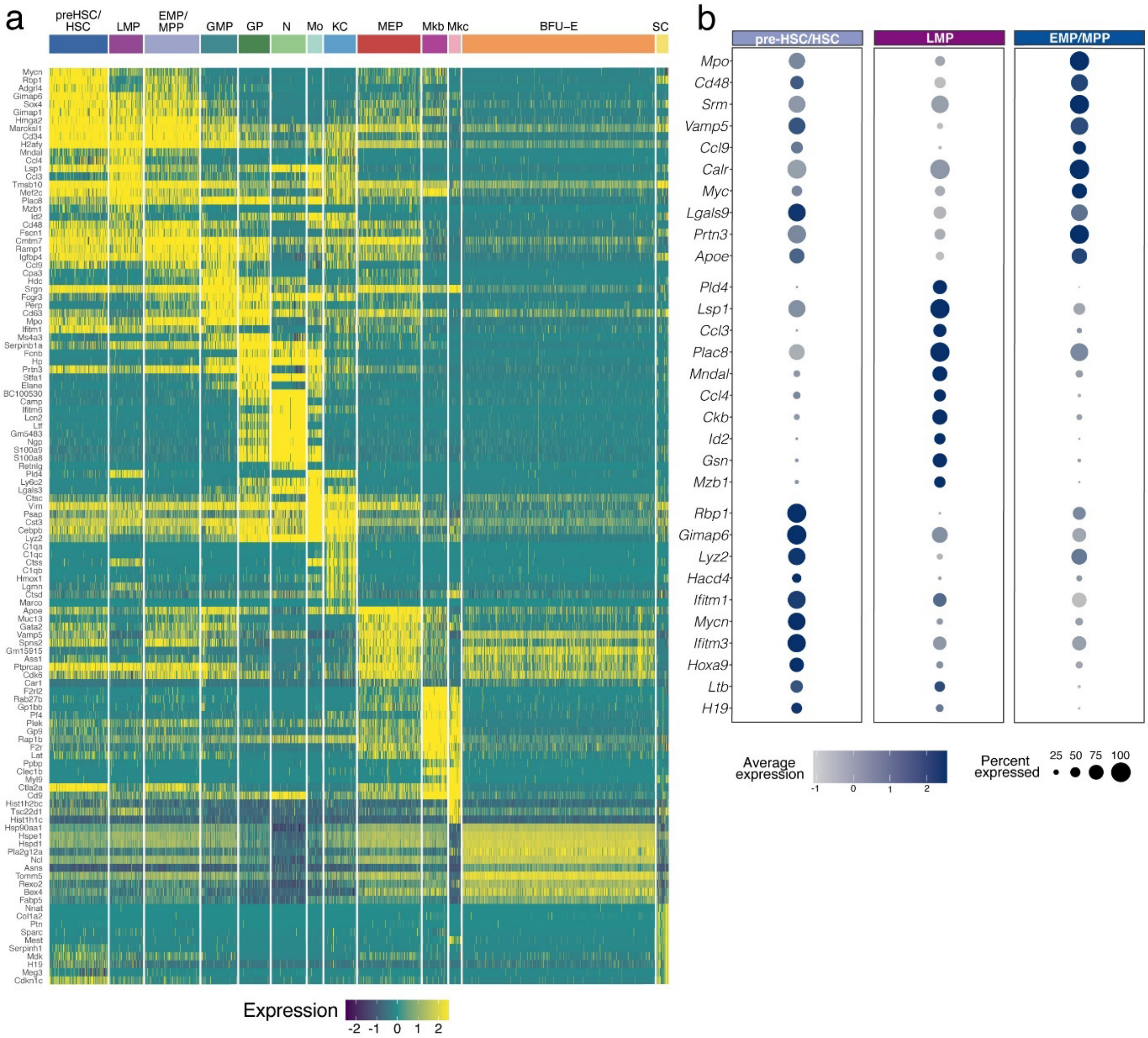


**Figure S2. Heatmap of differential gene expression in haematopoietic cell populations of the E12.5 mouse liver.**

(**a**) A heatmap plot shows DEGs among the haematopoietic cell clusters.

(**b**) A bubble plot shows DEGs among the haematopoietic progenitor subclusters.

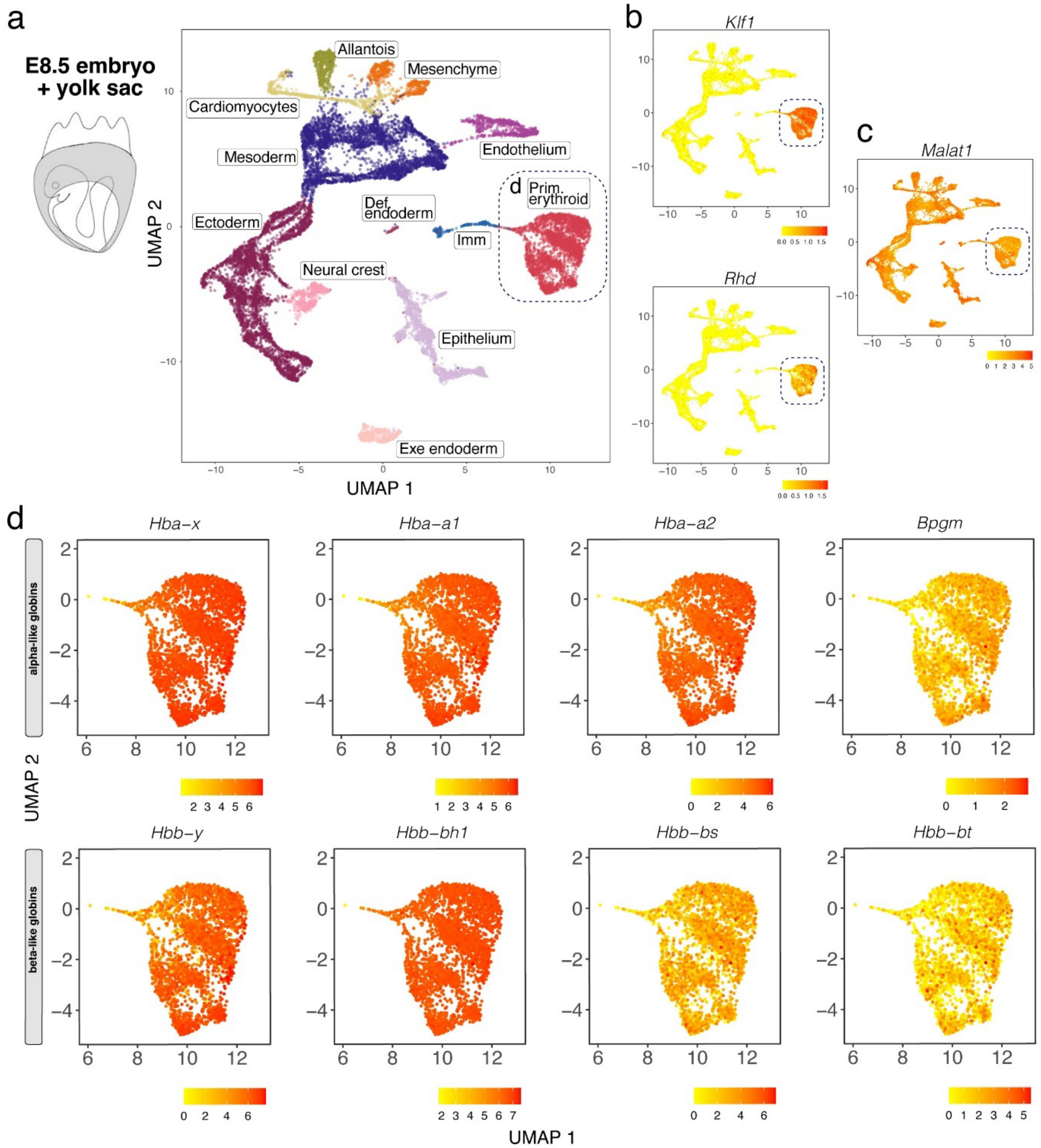


**Figure S3**. **scRNA-seq of E8.5 mouse embryos and their yolk sacs.**

(**a-c**) UMAP plots visualise clusters of distinct cell types (**a**), the expression of the indicated erythrocyte markers (**b**) and the nuclear marker *Malat1* (**c**) in the total E8.5 embryo and yolk sac; the stippled boxes indicate the erythroid cell subcluster shown in (**d**). (**d**) UMAP plots visualise the expression of the indicated haemoglobin genes and *Bpgm* in the erythroid cell subcluster.

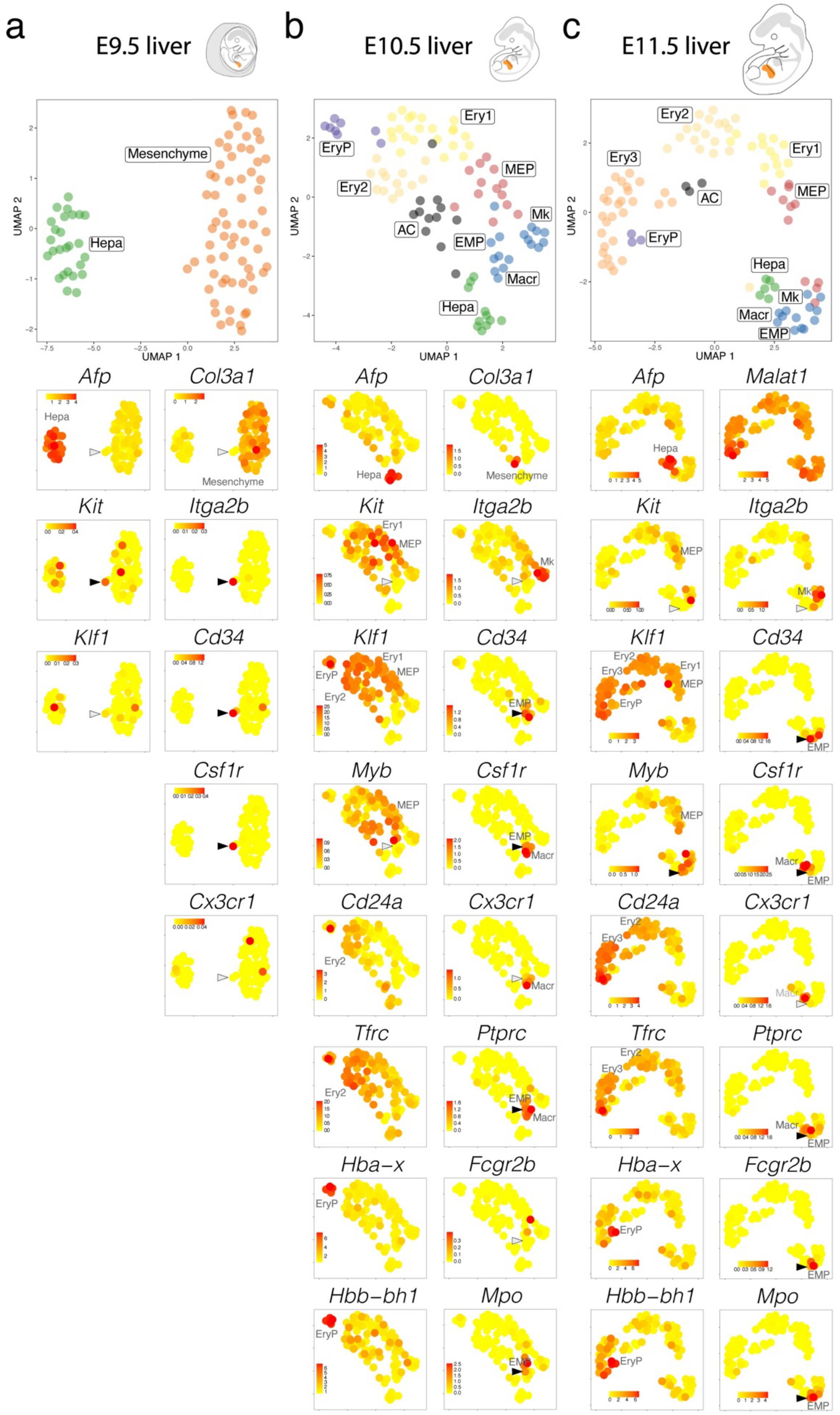


**Figure S4**. **scRNA-seq analysis of E9.5, E10.5 and E11.5 mouse livers.**
(**a-c**) scRNA-seq time-course of E9.5 (**a**), E10.5 (**b**) and E11.5 (**c**) mouse liver. UMAP plots visualise clusters of distinct cell types (**top panels**), and the expression of the indicated genes (**bottom panels**). AC, apoptotic cells; ND, not detected. Arrowheads and empty arrowheads point to EMPs/MPPs expressing or not the indicated gene, respectively. Each UMAP plot names the cluster(s) expressing the indicated gene.

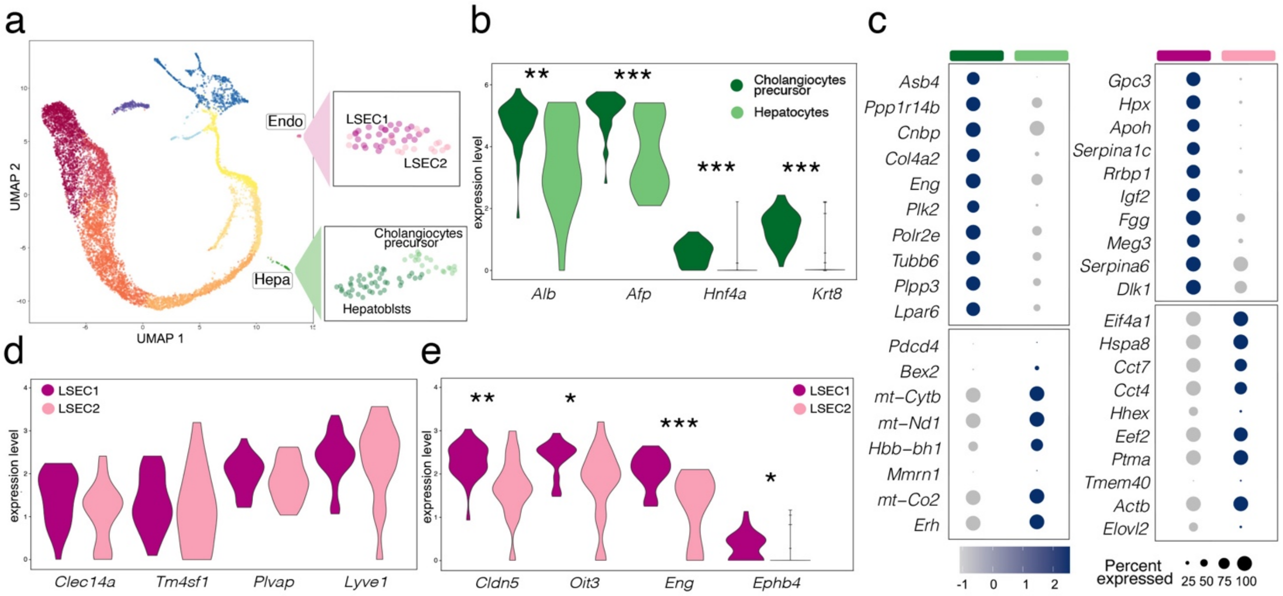


**Figure S5**. **Non-haematopoietic cell identification in E12.5 mouse liver by scRNA-seq analysis.**

(**a**) Subset selection followed by UMAP plot visualisation identifies subclusters of distinct non-hematopoietic cell types.

(**b-e**) Expression of the indicated genes in each subcluster, shown as violin plots (**b,d,e**) and bubble plots (**c**); * p>0.05, ** p<0.01, *** p<0.001 (non-parametric Wilcoxon rank sum test).
